# SARS-CoV-2 Spike Protein Impairs Endothelial Function via Downregulation of ACE2

**DOI:** 10.1101/2020.12.04.409144

**Authors:** Yuyang Lei, Jiao Zhang, Cara R. Schiavon, Ming He, Lili Chen, Hui Shen, Yichi Zhang, Qian Yin, Yoshitake Cho, Leonardo Andrade, Gerry S. Shadel, Mark Hepokoski, Ting Lei, Hongliang Wang, Jin Zhang, Jason X.-J. Yuan, Atul Malhotra, Uri Manor, Shengpeng Wang, Zu-Yi Yuan, John Y-J. Shyy

## Abstract

Coronavirus disease 2019 (COVID-19) includes the cardiovascular complications in addition to respiratory disease. SARS-CoV-2 infection impairs endothelial function and induces vascular inflammation, leading to endotheliitis. SARS-CoV-2 infection relies on the binding of Spike glycoprotein (S protein) to angiotensin converting enzyme 2 (ACE2) in the host cells. We show here that S protein alone can damage vascular endothelial cells (ECs) in vitro and in vivo, manifested by impaired mitochondrial function, decreased ACE2 expression and eNOS activity, and increased glycolysis. The underlying mechanism involves S protein downregulation of AMPK and upregulation of MDM2, causing ACE2 destabilization. Thus, the S protein-exerted vascular endothelial damage via ACE2 downregulation overrides the decreased virus infectivity.

---

Clinical data from coronavirus disease 2019 (COVID-19) underscore the cardiovascular complications in addition to respiratory disease involved in this deadly disease. SARS-CoV-2 infection relies on the binding of Spike glycoprotein (S protein) to angiotensin converting enzyme 2 (ACE2) in the host cells. Vascular endothelium is a key tissue infected and damaged by SARS-CoV-2.^1^ SARS-CoV-2 infection triggers mitochondrial ROS production and promotes glycolytic shift.^2^ Paradoxically, ACE2 exerts a protective effect on the cardiovascular system, and SARS-CoV-1 S protein has been shown to promote lung injury by decreasing the level of ACE2 in the infected lungs.^3^ In the current study, we show that S protein alone can damage vascular endothelial cells (ECs) by downregulating ACE2 and consequently inhibiting mitochondrial function.

To examine whether S protein causes lung damage in the infected host, we administered a pseudovirus expressing S protein (Pseu-Spike) to hamsters intratracheally. Lung damage was apparent in animals receiving Pseu-Spike, revealed by thickening of the alveolar septa and increased infiltration of mononuclear cells (Figure 1A). AMPK phosphorylates ACE2 Ser-680, MDM2 ubiquitinates ACE2 Lys-788, and crosstalk between AMPK and MDM2 determines the ACE2 expression level.^4^ In the damaged lungs, levels of phospho-AMPK (pAMPK), phospho-ACE2 (pACE2), and ACE2 decreased but those of MDM2 increased (Figure 1B*a*). Furthermore, complementary increased and decreased phosphorylation of eNOS Thr-494 and Ser-1176 indicated impaired eNOS activity. These changes of pACE2/ACE2 and MDM2 expression and AMPK activity in the endothelium by S protein were recapitulated by *in vitro* experiments using cultured pulmonary arterial ECs (PAECs) infected with Pseu-Spike which was rescued by treatment with N-acetyl-L-cysteine (NAC), the ROS inhibitor (Figure 1B*b*).

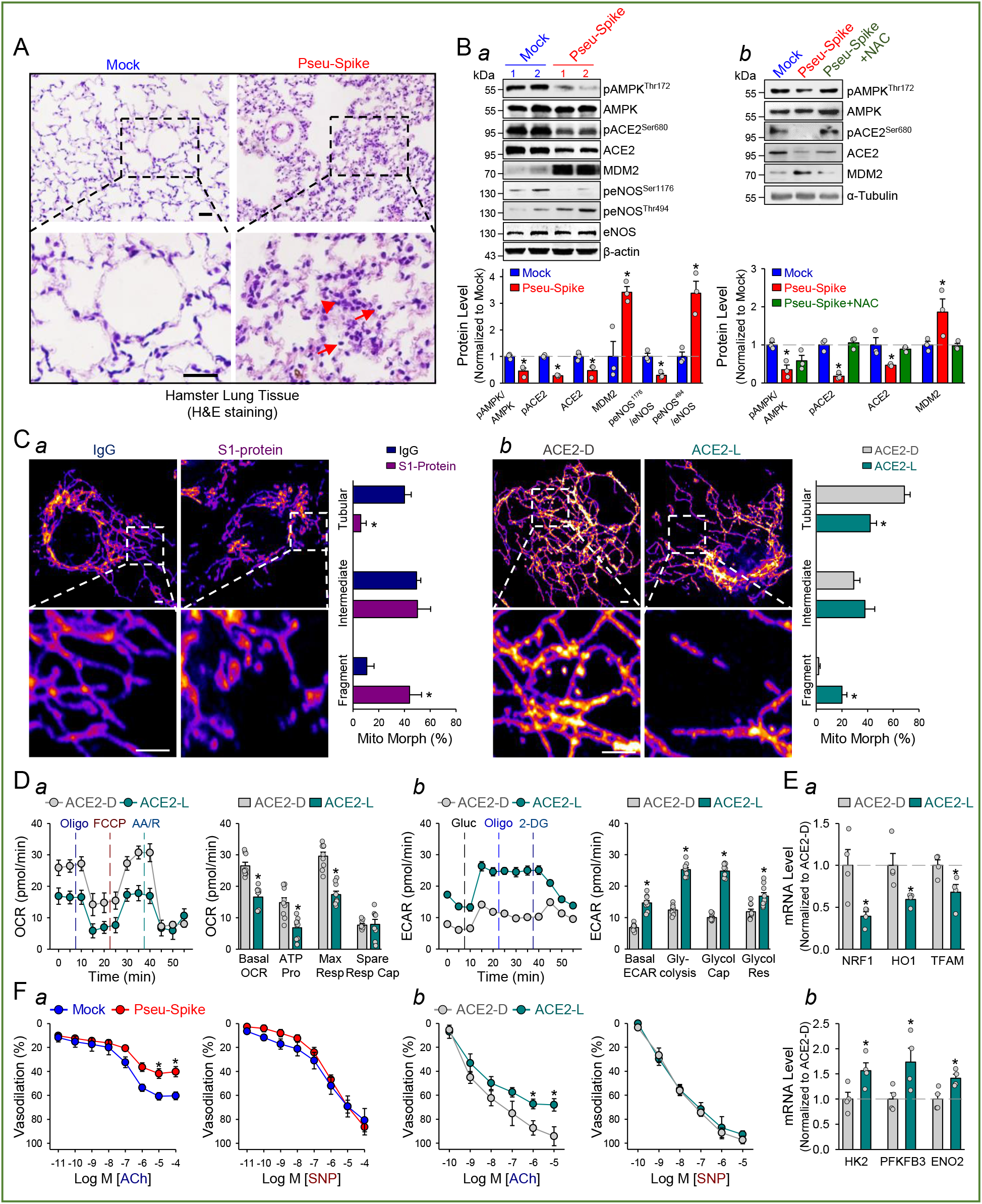
SARS-CoV-2 Spike protein exacerbates EC function via ACE2 downregulation and mitochondrial impairment. **A,** Representative H&E histopathology of lung specimens from 8-12 week-old male hamsters 5-day post administration of pseudovirus overexpressing Spike protein (Pseu-Spike) or those receiving mock virus in control group (n=3 mice per group, 5×10^8^ PFU). Thickened alveolar septa (red arrowhead) and mononuclear cell (red arrow). Scale bar = 20 μm. **B**, Lung tissues isolated from the Pseu-Spike (n=3) or mock virus (n=3)-infected hamsters were subjected to Western blot analysis for pAMPK T172, AMPK, pACE2 S680, ACE2, MDM2, p-eNOS S1176, p-eNOS T494, eNOS, and β-actin (**B***a*). Human pulmonary artery endothelial cells (PAECs) were infected with Pseu-Spike or mock virus for 24 hr with or without NAC (5 nmol/L) pre-treatment for 2 hr. The protein extracts were analyzed by Western blot using antibody for pAMPK T172, AMPK, pACE2 S680, ACE2, MDM2 and α-tubulin (n=3) (**B***b*). **C,** Representative confocal images of mitochondrial morphology of EC treated with recombinant S1 protein or human IgG (4 μg/ml) for 24 hr (**C***a*) or infected with human adenovirus ACE2 S680D (ACE2-D) or ACE2 S680L (ACE2-L) (10 MOI) for 48 hr (**C***b*). Mitochondria were visualized using TOM20 antibody (n=3, 50 cells counted for each replicate). Scale bar = 2.5 μm. **D,** Seahorse Flux assay of oxygen consumption rate (OCR, **D***a*) and extracellular acidification rate (ECAR, **D***b*) in ECs infected with ACE2-D or ACE2-L (10 MOI) for 48 hr (n=3). **E,** RT-qPCR analysis of the indicated mRNA levels in lung ECs from ACE2-D (n=4) and ACE2-L (n=4) knock-in mice. **F,** Dose-response curves of acetylcholine (ACh, left panels)- and sodium nitroprusside (SNP, right panels)-mediated relaxation on the tension of phenylephrine (1 μmol/L) precontracted intrapulmonary artery stripes from Pseu-Spike-(ACh n=8, SNP n=5) or mock-(ACh n=6, SNP n=5) virus infected hamsters (5 × 10^8^ PFU) (**F***a*) and ACE2-D (n=6) or ACE2-L (n=5) mice (**F***b*). The animal experiments were approved by the ethical committee of Xi’an Jiaotong University. Data are mean values ± SEM; *p <0.05 (two-tailed Student-*t* test in B*a*, C, D, E, F and one-way ANOVA in B*b*).

Since mitochondrial homeostasis has a profound effect on EC function, we next studied the impact of S protein on mitochondrial function. Double-blind analysis of confocal images of ECs treated with S1 protein revealed increased mitochondrial fragmentation, indicating altered mitochondrial dynamics (Figure 1C*a*). To examine whether these changes in mitochondrial morphology were due, in part, to the decreased amount of ACE2, we overexpressed ACE2 S680D (ACE2-D, a phospho-mimetic ACE2 with increased stability) or S680L (ACE2-L, a dephospho-mimetic with decreased stability)^4^ in ECs. As shown in Figure 1C*b*, ECs with ACE2-L overexpression had a higher number of fragmented mitochondria when compared to those with ACE2-D. Next, we performed Seahorse Flux analysis to compare the effect of ACE2-D versus ACE2-L on mitochondrial function. Seahorse Flux analysis showed ECs overexpressing ACE2-L had reduced basal mitochondrial respiration, ATP production, and maximal respiration compared to ECs overexpressing ACE2-D (Figure 1D*a*). Moreover, ACE2-L overexpression caused increased basal acidification rate, glucose-induced glycolysis, maximal glycolytic capacity, and glycolytic reserve (Figure 1D*b*). To explore the molecular basis underlying ACE2 regulation of mitochondrial and glycolytic alterations, we assessed the expression levels of mitochondria-and glycolysis-related genes in lung ECs isolated from ACE2-D or ACE2-L knock-in mice^4^. As shown in Figure 1E, the mRNA levels of NRF1, HO1, and TFAM (mitochondria biogenesis related genes) were increased, whereas those of HK2, PFKFB3, and ENO2 (glycolysis related genes) were decreased in the lung ECs in ACE2-D mice, as compared with those in ACE2-L mice.

SARS-CoV-2 infection impairs EC function and induces inflammation, leading to endotheliitis.^1,5^ Because S protein decreased the level of ACE2 and impaired eNOS derived-NO bioavailability, we examined whether S protein entry is indispensable for dysfunctional endothelium *in vivo*. As shown in Figure 1F*a*, the endothelium-dependent vasodilation induced by acetylcholine (ACh) was impaired in pulmonary arteries isolated from Pseu-Spike-administered hamsters, whereas the endothelium-independent vasodilation induced by sodium nitroprusside (SNP) was not affected by Pseu-Spike. To further confirm that EC-dependent vasodilation is correlated with ACE2 expression level, we compared the ACh-and SNP-induced vasodilation of pulmonary vessels from ACE2-D or ACE2-L mice. As anticipated, ACh-induced vasodilation was significantly hindered in pulmonary arteries isolated from ACE2-L mice in comparison to ACE2-D mice (Figure 1F*b*). There was, however, no difference of SNP-induced vasodilation in ACE2-D and ACE-L animals.

Our data herein reveals that S protein alone can damage endothelium, manifested by impaired mitochondrial function and eNOS activity but increased glycolysis. The S protein-mediated endothelial impairment depends on ACE2 post-translational modifications. It appears that S protein in ECs increases redox stress which may lead to AMPK deactivation, MDM2 upregulation, and ultimately ACE2 destabilization.^4^ It seems paradoxical that ACE2 reduction by S protein would decrease the virus infectivity, thereby protecting endothelium. However, a dysregulated renin-angiotensin system due to ACE2 reduction may exacerbate endothelial dysfunction, leading to endotheliitis. Collectively, our results suggest that the S protein-exerted EC damage overrides the decreased virus infectivity. This conclusion suggests that vaccination-generated antibody and/or exogenous antibody against S protein not only protects the host from SARS-CoV-2 infectivity but also inhibits S protein-imposed endothelial injury and ultimately decrease cardiovascular complication-associated mortality in COVID-19 patients.

## Data availability

The data that support the findings of this study are available from the corresponding author upon request.

